# On set-based association tests: insights from a regression using summary statistics

**DOI:** 10.1101/673103

**Authors:** Yanyan Zhao, Lei Sun

**Affiliations:** Department of Statistical Sciences, University of Toronto, 100 St. George Street, Toronto, Ontario M5S 3G3, Canada; Division of Biostatistics, Dalla Lana School of Public Health, University of Toronto, 155 College Street, Toronto, Ontario M5T 3M7, Canada

**Keywords:** Correlation, Summary statistics, Regression, Set-based tests, Sparse alternatives

## Abstract

Motivated by, but not limited to, association analyses of multiple genetic variants, we propose here a summary statistics-based regression framework. The proposed method requires only variant-specific summary statistics, and it unifies earlier methods based on individual-level data as special cases. The resulting score test statistic, derived from a linear mixed-effect regression model, inherently transforms the variant-specific statistics using the precision matrix to improve power for detecting sparse alternatives. Furthermore, the proposed method can incorporate additional variant-specific information with ease, facilitating omic-data integration. We study the asymptotic properties of the proposed tests under the null and alternatives, and we investigate efficient p-value calculation in finite samples. Finally, we provide supporting empirical evidence from extensive simulation studies and two applications.

---

Set-based joint analyses of multiple variables are increasingly important in many current scientific studies. For example, in modern genome-wide association studies (GWAS), one might be interested in jointly analyzing multiple (rare) genetic variants influencing a complex, and heritable trait (also known as gene- or pathway-based association studies), identifying one genetic variant influencing multiple traits (also known as pleitrotropy studies), or combining evidence from multiple studies as in the classical meta-analyses.

Without loss of generality, let us focus on set-based analyses of multiple rare genetic variants. In this setting, myriad statistical tests have been proposed and they fit into three general categories (e.g. Derkach, Lawless & Sun, 2014; Lee et al., 2014): the linear or burden tests (e.g. Madsen & Browning, 2009), the quadratic or variance-component tests (e.g. Pan, 2009; Wu & Lin, 2011), and the hybrid tests combining evidence from the linear and quadratic tests (e.g. Lee, Wu & Lin, 2012; Derkach, Lawless & Sun, 2013).

These earlier methods have been studied extensively, but they can be improved in several aspects. First, these methods may not perform well in the sparse-signal setting where only a small proportion of the variants are truly associated (Xu et al., 2016). Second, individual/subject-level data may not be available in practice, so it is useful to develop summary statistics-based association tests, and ideally the tests can also incorporate additional information available for each variant. Finally, earlier work have shown that the performance of a simple minimum-p value approach (Derkach, Lawless & Sun, 2013) is comparable with that of the optimal sequence kernel association test, SKAT-O (Lee, Wu & Lin, 2012). This suggests that a grid search for the ‘optimal’ weighting factor may not be necessary, and it is helpful to develop new robust hybrid test statistics with model-driven weights.

To this end, we propose a flexible and unifying linear mixed-effect regression model that requires only variant-specific summary statistics, and we show that earlier methods based on individual-level data are special cases of the proposed testing framework. The set-based association test statistic derived from the regression model inherently transforms the variant-specific summary statistics using the precision matrix to improve power for detecting sparse alternatives; see Fan, Jin & Yao (2013) and Cai, Liu & Xia (2014) for the use of precision matrix in other high-dimension analytical settings. Furthermore, the proposed method can incorporate additional variant-specific information as a covariate, e.g. the functional importance of the variants to be analyzed (Ionita-Laza et al., 2016). Through simulation and application studies, we show that the proposed method can have substantial power gain when the included covariate contains useful information, while power loss is minimal when the covariate is uninformative.

Although the proposed set-based regression approach is motivated by joint analyses of multiple rare genetic variants, the method can be used in other settings, for example, meta-analyses of multiple studies (Han & Eskin, 2011), data integration of GWAS and gene-expression data (Gamazon et al., 2015), pleiotropy association studies of multiple phenotypes (Liu & Lin, 2018b), as well as polygenic risk score analyses of multiple common variants (Purcell et al., 2009). We will discuss later the differences and connections between the proposed method and these earlier work.

The remainder of the paper is organized as follows. Section 1 reviews existing association methods for jointly analyzing a set of rare genetic variants based on individual-level data. Section 2 first outlines the proposed regression framework based on summary statistics, from which we derive a catalog of association test statistics from fixed-, random- or mixed-effect models. We then demonstrate the analytical equivalence between some of the newly proposed statistics and the existing ones for rare variants analyses. This section also investigates efficient p-value calculation in finite samples, studies the asymptotic properties of the proposed tests, and considers covariate adjustments. Section 3 provides supporting empirical evidence from extensive simulation studies, and Section 4 presents results from two application studies. Section 5 concludes with remarks and discussion, and the Supplementary Material provides theoretical proofs and additional numerical results.

## 1 Existing association tests for jointly analyzing a set of rare genetic variants

### 1.1 Regression set-up using individual-level data

Consider a sample of *n* independent individuals and a set of *J* genetic variants of interest, let ***y*** = (*y*_1_, …, *y*_*n*_)^*T*^ denote the phenotype variable and ***G*** be a *n* × *J* matrix for the corresponding genotypes with elements *G*_*i j*_, *i* = 1, …, *n* and *j* = 1, …, *J*, ***G***_*i*_ = (*G*_*i*1_, *G*_*i*2_, …, *G*_*iJ*_)^*T*^ and ***G*** _*j*_ = (*G*_1 *j*_, *G*_2 *j*_, …, *G*_*n j*_)^*T*^. Assume that *y*_*i*_ given ***G***_*i*_ follows an exponential family distribution with mean *µ*_*i*_ and consider using the canonical link function *g*(·) and

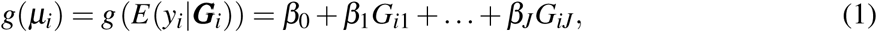

to model the phenotype-genotype association, where *β* _*j*_ is the regression coefficient for genetic variant *j*. Individual-level covariates information such as age and sex, if available, should be added to the model but are omitted for the moment for notation simplicity.

We are interested in testing the hypothesis that

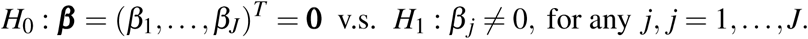

The corresponding score vector is

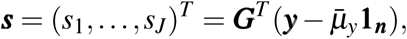

where **1**_***n***_ is a *n* × 1 unit vector, 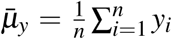 and 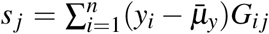 capturing the linear relationship between phenotype ***y*** and genotype ***G*** _*j*_. The variance-covariance matrix of ***s*** is

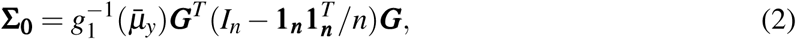

where *g*_1_(·) denotes the first derivative of the link function *g*(·), and *I*_*n*_ is a identity matrix of size *n*.

### 1.2 Existing methods based on the score vector *s*

Although it may not be obvious, most test statistics developed for rare variants analyses are functions of ***s***. For example, the original burden test (Madsen & Browning, 2009) first constructs a ‘super-allele’, 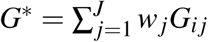, where ***w*** = (*w*_1_, *w*_2_, …, *w*_*J*_)^*T*^ are pre-specified weights often associated with the minor allele frequency (MAF) of the variants. The burden test then considers regression model, 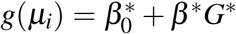 and test *H*_0_ : *β*^*^ = 0, v.s. *H*_1_ : *β*^*^ ≠ 0. However, it is not difficult to show that the score test statistic derived from the regression using *G*^*^ is proportional to

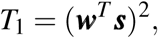

where under the null of no association (***w***^**′**^ **Σ**_**0**_ ***w***)^−1^ *T*_1_ follows a central chi-square distribution with 1 degrees of freedom (d.f.), 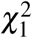, for a fixed *J* but assuming that ***s*** is multivariate normal asymptotically with respect to *n* (Derkach, Lawless & Sun, 2014).

This *T*_1_ test is also termed as the sum test by Pan (2009), and it belongs to the linear class of tests (Derkach, Lawless & Sun, 2014). Because *T*_1_ is based on the weighted average of *s* _*j*_, and *s* _*j*_ can be positive or negative depending on the direction of effect (i.e. sign of *β* _*j*_ in model (1)), *T*_1_ is only powerful when a large proportion of variants are causal *and* effects are in the same direction. Variance-component tests, such as SSU (Pan, 2009) and SKAT (Wu & Lin, 2011), are alternatives that belong to the quadratic class of tests; see Table 1 of Derkach, Lawless & Sun (2014)for a summary. Again, although most of these tests started with regression model (1), they can be formulated as

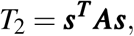

where 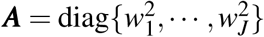 and *w*_*j*_ depends on the MAF of variant *j, j* = 1, …, *J*. Under the null and the multivariate normality assumption for ***s***, *T*_2_ follows a weighted sum of 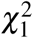 distribution for a fixed *J* (Derkach, Lawless & Sun, 2014). *T*_2_ is robust to heterogenous effect directions, but it is less powerful than *T*_1_ when most variants are causal and with the same direction of effects.

**Table 1:**
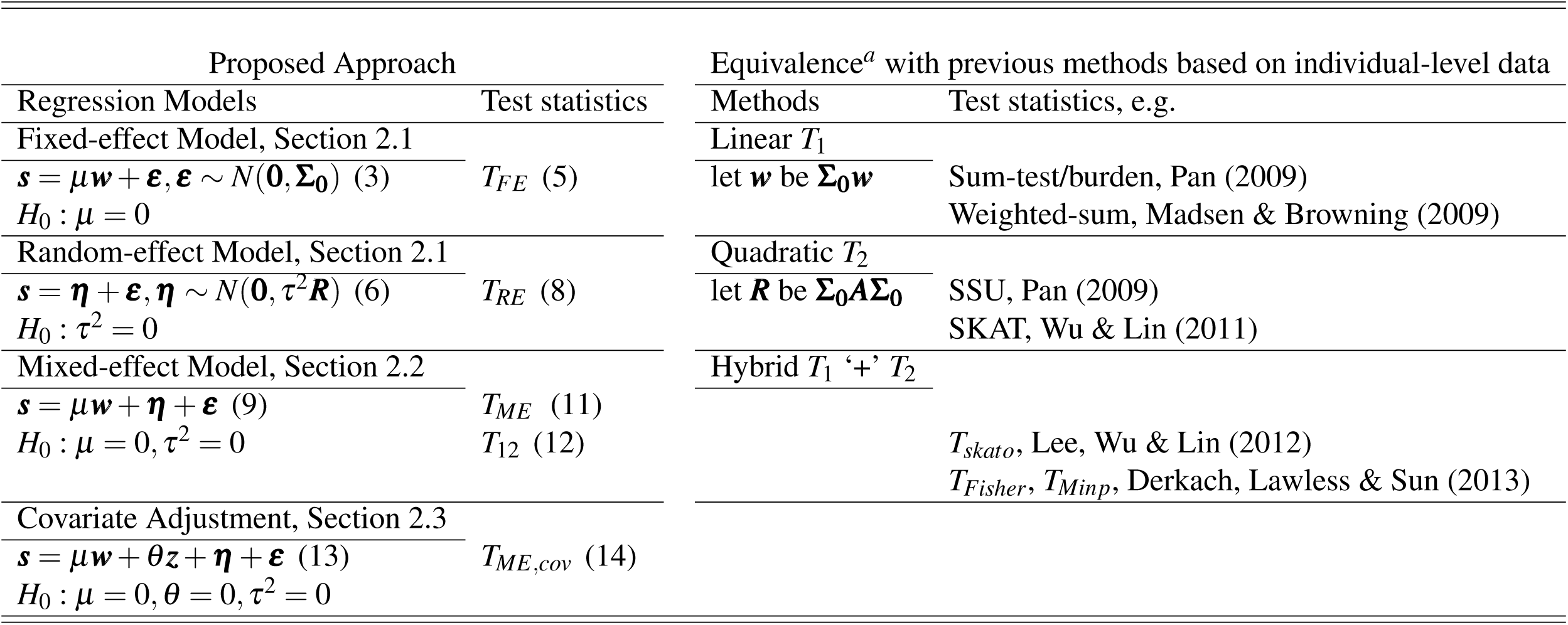
Summary of different test statistics for analyzing a set of genetic variants. ^*a*^The equivalence can be analytical or numerical; see corresponding sessions for details. See Table 1 of Derkach, Lawless & Sun (2014) for a more detailed summary of the previous methods.

Since the true genetic model is unknown in practice, hybrid tests combining *T*_1_ and *T*_2_ have been proposed. For example, SKAT-O of Lee, Wu & Lin (2012) considers

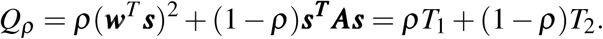

It then performs a grid search for the ‘optimal’ *ρ*, 0 = *ρ*_1_ < *ρ*_2_ < … < *ρ*_*m*_ = 1. Let *p*_*ρ*_ be the p-value corresponding to each *Q*_*ρ*_, the final SKAT-O test statistic is

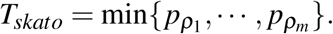

The asymptotic p-value of *T*_*skato*_ can be calculated with one-dimensional numerical integration (Lee, Wu & Lin, 2012).

Instead of considering data-driven ‘optimal’ *ρ* and adjusting for the inherent selection bias, Derkach, Lawless & Sun (2013) proposed two simpler yet competitive hybrid test statistics, *T*_*Fisher*_ and *T*_*Minp*_. Let 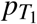 and 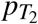 be the p-values corresponding to *T*_1_ and *T*_2_, respectively, the Fisher and Minp statistics take the form of

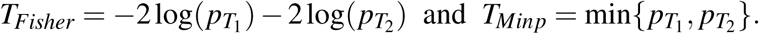

Under the null of no association, 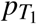 and 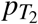 are asymptotically independent of each other, thus *T*_*Fisher*_ is 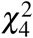 distributed and *T*_*Minp*_ is *Beta*(1, 2) distributed.

Derkach, Lawless & Sun (2013) have also shown that *T*_*Minp*_ and *T*_*skato*_ perform similarly, and they are slightly more powerful than *T*_*Fisher*_ when *T*_1_ has no power (e.g. less than 20%). On the other hand, when both *T*_1_ and *T*_2_ have some power *T*_*Fisher*_ is more powerful than *T*_*Minp*_ and *T*_*skato*_. However, we expect all three hybrid tests to have diminished power of detecting sparse alternatives (Donoho & Jin, 2004; Barnett, Mukherjee & Lin, 2017) when only a small proportion of the variants in a set are truly associated.

To improve performance, we first note that if variants are correlated with each other, we can consider for example 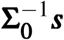 instead of ***s*** to construct a more powerful test for detecting sparse alternatives. Second, it is clear now that we only need variant-specific summary statistics ***s*** = (*s*_1_, …, *s*_*J*_)^*T*^ to joint analyze the *J* variants of interest, provided **Σ**_**0**_ can be accurately estimated. Further, the fact that *T*_*Minp*_ and *T*_*skato*_ having similar performance suggests that a grid search for *ρ* may not be necessary, and an easy-to-compute yet theoretically justified ‘optimal’ *ρ* may exist. Lastly, when additional variant-specific information *z* _*j*_ (e.g. variant *j* being non-synonymous or not) is available, we can improve power by incorporating ***z*** = (*z*_1_, …, *z*_*J*_)^*T*^. Intuitively we can achieve this by modifying *w*_*j*_ to incorporate *z* _*j*_ in additional to the MAF, but a less add-hoc approach is desirable. In the section, we develop a flexible and unifying regression framework that(i) requires only *s* _*j*_ and **Σ**_**0**_, (ii) yields *T*_1_ and *T*_2_ as special cases and a hybrid statistic similar to *T*_*skato*_, and (iii) provides new test statistics that incorporate the precision matrix 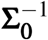 and account for covariate information ***z***.

## 2 A regression framework based on summary statistics

Here we assume that ***s*** = (*s*_1_, …, *s*_*J*_)^*T*^ is available, summarizing the association relationship between the phenotype of interest and a set of *J* genetic variants as detailed in Section 1. We also assume that **Σ**_**0**_ is known or estimated accurately from a reference panel (Cheng et al., 2019; Fan, Liao & Liu, 2016). Let ***z*** = (*z*_1_, …, *z*_*J*_)^*T*^ represent variant-specific information available.

### 2.1 Fixed-effect and random-effect models

We first consider a fixed-effect (FE) model that models the common effect *µ* present among *s* _*j*_, *j* = 1, …, *J*,

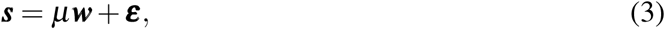

where ***w*** = (*w*_1_, …, *w*_*J*_)^*T*^, ***ε*** = (*ε*_1_, …, *ε*_*J*_)^*T*^ and ***ε*** ∼ *N*(**0, Σ**_**0**_). Based on this model, we test

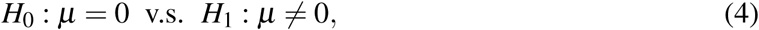

and the corresponding score test statistic is

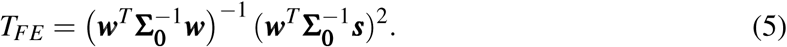

If we let *w* in *T*_*FE*_ to be **Σ**_**0**_***w***, the analytical equivalence between *T*_*FE*_ and *T*_1_ in Section 1 is easy to show because *T*_1_ ∝ (***w***^*T*^ **Σ**_**0**_***w***)^−1^ (***w***^*T*^ ***s***)^2^ (Table 1).

Alternatively, we can consider the following random-effect (RE) model,

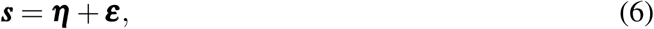

where ***η*** ∼ *N*(**0**, *τ*^2^ ***R***), ***R*** is a *J* × *J* pre-defined positive or semi-definite symmetric matrix, and ***ε*** as defined above. If we test

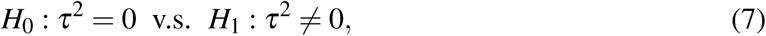

the corresponding score test statistic is

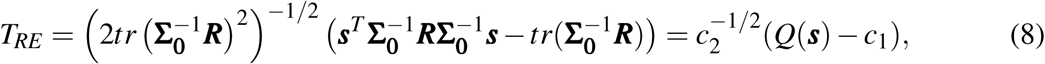

where 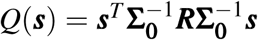, and 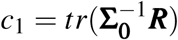 and 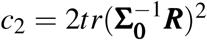 are respectively the mean and variance of *Q*(***s***). If we let ***R*** = **Σ**_**0**_***A*Σ**_**0**_, the analytical equivalence between *T*_*RE*_ and *T*_2_ = ***s***^*T*^ ***As*** is also apparent because *T*_*RE*_ ∝ *Q*(***s***) (Table 1). In addition, if we let ***R*** = **Σ**_**0**_***W R***_***ρ***_ ***W* Σ**_**0**_, where ***W*** = diag{*w*_*j*_} and 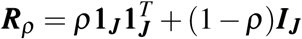, then *Q*(***s***) = *ρT*_1_ + (1 −*ρ*)*T*_2_ = *Q*_*ρ*_, the key element of *T*_*skato*_.

### 2.2 Mixed-effect model

The fixed-effect model (3) captures the common underlying effect, while the random-effect model (6) accounts for potential effect heterogeneity. A logical next step is to consider a mixed-effect (ME) modelling framework that includes models (3) and (6) as special cases,

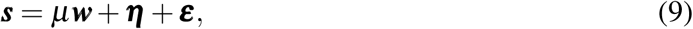

where ***w, η*** and ***ε*** are defined as before. If we test

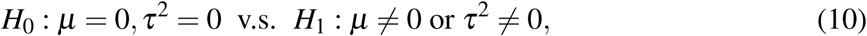

the corresponding score vector is 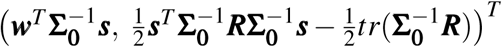, and the test statistic is

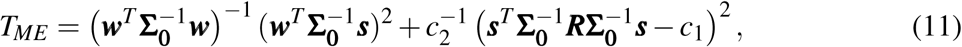

where 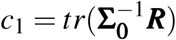 and 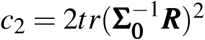, the same as for *T*_*RE*_.

Note that *T*_*ME*_ uses 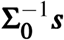 to account for the correlation between the tested variants. Transforming ***s*** by the precision matrix has been considered previously in other settings. For example, Cai, Liu & Xia (2014) showed that the transformation can improve power of a two-sample high-dimensional means test for detecting sparse alternatives in the presence of high correlation.

Following the construction of *T*_*ME*_, intuitively we can also consider the following weighted average of *T*_1_ = (***w***^*T*^ ***s***)^2^ and *T*_2_ = ***s***^*T*^ ***As*** (and ***A*** = ***R***),

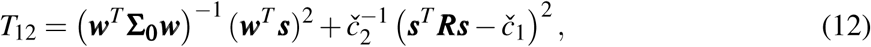

where *č*_1_ = *tr*(**Σ**_**0**_***R***) and *č*_2_ = 2*tr*(**Σ**_**0**_***R***)^2^ are respectively the mean and variance of *T*_2_ under the *H*_0_ of (10). The connection between *T*_12_ with *T*_*skato*_ is also immediate. However, there are two key differences. First, given a *ρ, T*_*skato*_ relies on *Q*_*ρ*_ = *ρT*_1_ + (1 − *ρ*)*T*_2_. In contrast, *T*_12_ combines *T*_1_ and the *square* of *centralized T*_2_ (not *T*_2_ itself). Secondly, *T*_*skato*_ searches for the ‘optimal’ *ρ* that minimizes the p-value associated with *Q*_*ρ*_ and then adjusts for selection bias. In contrast, *T*_12_ uses the variances of *T*_1_ and *T*_2_ as weights.

### 2.3 Covariate adjustment

Given ***z*** = (*z*_1_, …, *z*_*J*_)^*T*^, one could consider directly modifying ***w*** to reflect the additional information available for the *J* variants of interest, but a principled approach is lacking. The proposed regression framework, however, can naturally incorporate ***z*** as a covariate into the regression,

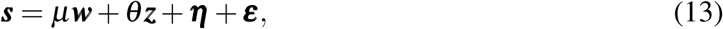

where ***w, η*** and ***ε*** are defined as before. If we are interested in testing,

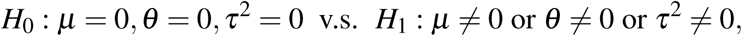

the corresponding score test statistic is

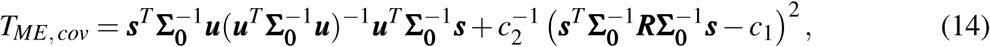

where ***u*** = (***w, z***), and *c*_1_ and *c*_2_ are defined as before. Table 1 summarizes all the tests discussed so far.

### 2.4 Asymptotic distributions of the proposed test statistics

The asymptotic distributions of the existing tests, *T*_1_, *T*_2_, *T*_*skato*_, *T*_*Fisher*_, and *T*_*Minp*_, have been previously established. Here, we establish the asymptotic distribution of *T*_*ME*_ and *T*_12_ when *J* → ∞. In the next section, we will study finite sample behaviour of the tests. We begin with some notations and mild conditions needed for Theorem 1. Let ***C*** be a matrix such that ***C*Σ**_**0**_***C***^*T*^ = ***I***_***J***_, and let *λ*_*min*_(·) and *λ*_*max*_(·) be the minimum and maximum eigenvalues of a matrix, respectively. For two sequences of real numbers {*a*_1*J*_} and {*a*_2*J*_}, denote *a*_1*J*_ = *o*(*a*_2*J*_) if lim_*J*→∞_(*a*_1*J*_*/a*_2*J*_) = 0.

**Condition 1.** 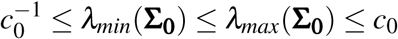 *for some constant c*_0_.

**Condition 2.** *c*^−1^ ≤ *λ*_*min*_(***R***) ≤ *λ*_*max*_(***R***) ≤ *c for some constant c.*

**Condition 3.** 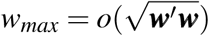 *where w*_*max*_ = max _*j*_ *w*_*j*_.

#### Theorem 1.

*Theorem 1.Under the null hypothesis of (10) and assume Conditions 1-3 hold*,

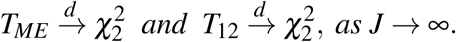

See the Supplementary Material for the proofs of Theorem 1 and all other theorems. When *J* is small, significance evaluation based on the asymptotic distributions may not be adequate. Theorem 2 provides an approximation for the finite-sample distribution of *T*_*ME*_; the result for *T*_12_ is similar.

#### Theorem 2.

*Theorem 2.Let λ* _*j*_ *be the jth eigenvalue of* ***CRC***^*T*^, *j* = 1, …, *J, then*

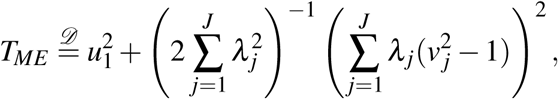

*where u*_1_ *and v* _*j*_, *j* = 1, …, *r, are independent N*(0, 1) *random variables, and* 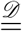 *denotes equality in distribution.*

We note that the above finite and asymptotic results are with respect to *J*. The validities of *T*_*ME*_ and *T*_12_ do not require *n* → ∞ explicitly, as long as ***s*** is multivariate normal.

### 2.5 Power comparison

In this section, we first establish the asymptotic distributions of *T*_*ME*_ and *T*_12_ (as *J* → ∞) under alternatives.

#### Theorem 3.

*Theorem 3.Under the alternative hypothesis* 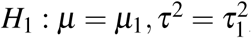,

a. *Assume Conditions 1-3 hold, then*

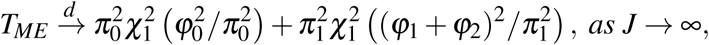

*where* 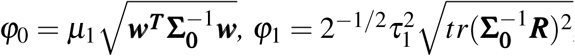, 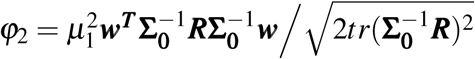, 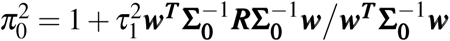, *and* 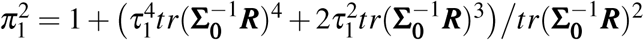.
b. *Assume Conditions 1-3 hold, then*

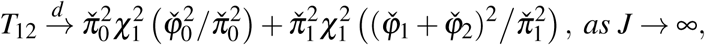

*where* 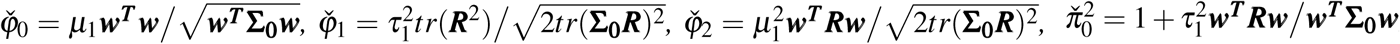, *and*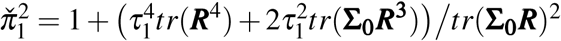.

To compare the asymptotic power between *T*_*ME*_ and *T*_12_, first let us consider the simple case of no random effect, i.e. 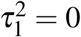. In that case, 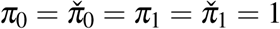, and we can show that *T*_*ME*_ is at least as powerful as *T*_12_ provided that 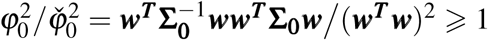. In fact, this inequality holds as long as **Σ**_**0**_ is a positive definite symmetric matrix. This suggests that if the true underlying model for ***s*** is a fixed-effect model, then *T*_*ME*_ is more powerful than *T*_12_. Our analytical conclusion here is consistent with that observed by Liu & Lin (2018a) for joint analyses of multiple phenotypes and by Brockwell & Gordon (2001) for meta-analyses.

We next consider a local alternative assuming

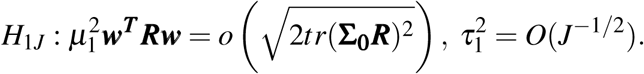

In this case, 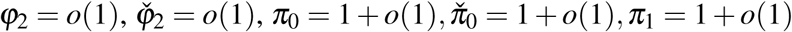, and 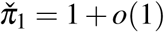 based on Conditions 1-2. As a result, *T*_*ME*_ is at least as powerful as *T*_12_, provided that 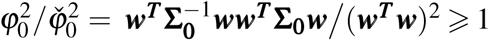 and 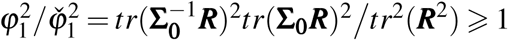. These two inequalities always hold as long as ***R*** and **Σ**_**0**_ are positive definite symmetric matrices.

## 3 Simulation studies

To compare the finite-sample performance of *T*_*ME*_ and *T*_12_, with *T*_*skato*_, *T*_*Minp*_ and *T*_*Fisher*_, we conduct extensive simulation studies, examining the effects of different correlation structures and signal sparsities on power.

To obtain the summary statistics, ***s*** and **Σ**_**0**_, for association studies of rare variants, similar to Derkach, Lawless & Sun (2014) we assume that the underlying individual-level model is *E*(*y*_*i*_|***G***_*i*_) = *β*_0_ + *β*_1_*G*_*i*1_ + … + *β*_*J*_*G*_*iJ*_, where *y*_*i*_ is normally distributed with variance *σ*^2^ = 1, *G*_*i j*_ is Bernoulli with *Pr*(*G*_*i j*_ = 1) = *p* _*j*_ where *p* _*j*_ is approximately twice the minor allele frequency of variant *j*, and *i* = 1, …, *n* and *j* = 1, …, *J*. Given this set-up, ***s*** ∼ *N*(***µ*, Σ**_**0**_), where 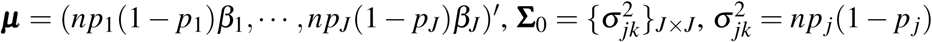 for *j* = *k* and 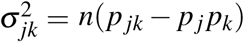 for *j* ≠ *k*, and *p*_*jk*_ = *P*(*G*_*i j*_ = 1, *G*_*ik*_ = 1).

We consider *n* = 500 and *J* = 10, 50, 100, 500, and 1000, and we draw *p* _*j*_ randomly from Unif (0.005, 0.02), *j* = 1, …, *J*. For 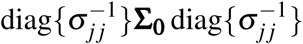, we consider an AR(1) pattern with correlation 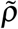, and 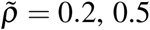 and 0.8. For ***w*** and ***A*** in *T*_*skato*_, *T*_*Minp*_, *T*_*Fisher*_, and *T*_12_, we choose the commonly used 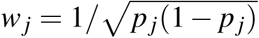 and 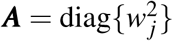. For ***w*** and ***R*** in *T*_*ME*_, we choose the same *w*_*j*_ and let ***R*** = ***A*** for a fair comparison.

### 3.1 Type I error

To examine the validity of the proposed tests, *T*_*ME*_ and *T*_12_, we generate ***s*** from *N*(**0, Σ**_**0**_), independently, 10^6^ times for each *J* and 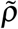 combination. The results in Table 2 show that for small *J*, in combination with stringent *α* level, the asymptotic p-values are not adequate. In that case, the approximate solution in Theorem 2 should be used.

**Table 2:** Type I error evaluation. Empirical test sizes for *α* = 5%, 1%, 0.1%, and 0.01%, estimated based on 10^6^ replications independently simulated under the null. *T*_.,*asy*_ represents p-value calculation using the asymptotic distributions in Theorem 1, and *T*_.,*apr*_ is for p-value estimation based on Theorem 2 with 10^7^ independent *u*_1_ and *v* _*j*_ standard normal variables generated for each of the 10^6^ simulated null replicates. Results here are for 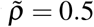; results for other 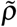 values are provided in the Supplementary Material.

| $J$ | $\alpha$ | $T_{12,asy}$ | $T_{me,asy}$ | $T_{12,apr}$ | $T_{me,apr}$ |
| --- | --- | --- | --- | --- | --- |
| 10 | 5% | 6.1558 | 5.6556 | 5.7470 | 5.2226 |
|  | 1% | 3.1219 | 2.4281 | 1.7649 | 1.2969 |
|  | 0.1% | 1.4531 | 0.9427 | 0.2079 | 0.1316 |
|  | 0.01% | 0.7725 | 0.4294 | 0.0196 | 0.0141 |
| 20 | 5% | 5.6845 | 5.5795 | 5.4308 | 5.2198 |
|  | 1% | 2.5849 | 2.4312 | 1.6450 | 1.3667 |
|  | 0.1% | 1.0219 | 0.9643 | 0.2212 | 0.1476 |
|  | 0.01% | 0.4703 | 0.4550 | 0.0246 | 0.0149 |
| 50 | 5% | 5.1085 | 5.1317 | 5.0333 | 4.9875 |
|  | 1% | 1.9088 | 1.7082 | 1.4498 | 1.1475 |
|  | 0.1% | 0.5952 | 0.5002 | 0.2103 | 0.1307 |
|  | 0.01% | 0.2148 | 0.1794 | 0.0230 | 0.0132 |
| 100 | 5% | 4.9508 | 5.0064 | 4.923 | 4.9664 |
|  | 1% | 1.5279 | 1.3854 | 1.2788 | 1.1030 |
|  | 0.1% | 0.3796 | 0.3082 | 0.1846 | 0.1273 |
|  | 0.01% | 0.1149 | 0.0831 | 0.025 | 0.0138 |
| 500 | 5% | 4.9335 | 5.0020 | 4.9307 | 4.9948 |
|  | 1% | 1.1265 | 1.0917 | 1.0720 | 1.0238 |
|  | 0.1% | 0.1662 | 0.1547 | 0.1288 | 0.1130 |
|  | 0.01% | 0.0332 | 0.0281 | 0.0202 | 0.0139 |
| 1000 | 5% | 4.9841 | 4.9552 | 4.9743 | 4.9436 |
|  | 1% | 1.0475 | 1.0254 | 1.0151 | 0.9864 |
|  | 0.1% | 0.1365 | 0.1190 | 0.1183 | 0.0951 |
|  | 0.01% | 0.0205 | 0.0162 | 0.0163 | 0.0109 |

For the existing methods, *T*_*skato*_, *T*_*Minp*_ and *T*_*Fisher*_, we observe in our simulation studies that *T*_*skato*_ is slightly conservative for the *α* levels considered regardless of the size of *J. T*_*Minp*_ is also slightly conservative for small *J* but has correct test size when *J* > 50, and *T*_*Fisher*_ has inflated type I error when the correlation among variants is strong (Supplementary Material). Our observations here are largely consistent with previous reports (e.g. Larson et al., 2017).

### 3.2 Power without covariates

We consider two different simulation designs to evaluate power. For both designs, *P*_*c*_, the proportion of causal variants for a given set of *J* variants, is randomly drawn from Unif(0.01, 0.1) for the case of sparse signals, and *P*_*c*_ ∼ Unif(0.1, 0.5) for the case of moderately sparse signals. Among the causal variants, *p*_*d*_ is the proportion of deleterious variants with *β* _*j*_ > 0 and *P*_*d*_ ∼ Unif (0.5, 0.75). We note that for each *P*_*c*_ and *P*_*d*_ combination, the locations of the signals (*β* _*j*_ ≠ 0) are randomly drawn from {1, 2, …, *J*}, without replacement. This randomness helps us to comprehensively explore the effect of different correlation structures between causal variants, between non-causal variants, as well as between causal and non-causal variants on power.

Design one follows the approach of Derkach, Lawless & Sun (2014). That is, ***s*** ∼ *N*(***µ*, Σ**_0_) and |*β* _*j*_| ∼ Unif(0.5, 1.5) for both the sparse and moderately sparse cases. Design two assumes that ***s*** is drawn from the mixed-effect model (9) with varying magnitudes of *τ*^2^. Specifically, for a given *J* we first simulate *p* _*j*_ ∼ Unif(0.005, 0.02), *j* = 1,*… J, P*_*c*_ ∼ Unif(0.1, 0.5) and *P*_*d*_ ∼ Unif(0.5, 0.75), and we draw the locations of the causal variants from {1, 2, …, *J*} as in design one. We then specify sign(*β* _*j*_) accordingly and obtain ***w***^**\***^ = (*np*_1_(1 − *p*_1_)sign(*β*_1_), …, *np*_*J*_(1 − *p*_*J*_)sign(*β*_*J*_))′; for a null variant sign(*β* _*j*_) = 0. Finally, we simulate ***s*** ∼ *N*(*β* ***w***^**\***^, *τ*^2^ ***I***_*J*_ + **Σ**_0_), where *β* ∼ Unif(0.5, 1.5) for both the sparse and moderately sparse cases, and *τ*^2^ ∼ Unif (1, 2).

For both designs, results of power comparison focus on *J* = 100, *α* = 0.05 and the sparse alternatives; results for the moderately sparse alternatives are characteristically similar (Supplementary Material). The empirical power for *α* = 0.05 are estimated from 10^3^ simulated replicates, and using the empirical critical values obtained from 10^4^ null replicates.

Figures 1(a) and 1(b) show that when the correlation among variants is relatively strong (e.g. 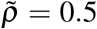 in Figures 1(a) and 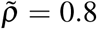 in the Supplementary Material), *T*_*ME*_ (based on 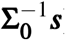) has better power than the other methods (based on ***s***) including *T*_*skato*_. However, this approach may not be advantageous when there is only weak correlation in conjunction with sparse signal (e.g. 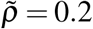 in Figures 1(b)) as discussed in Section 2.2. Interestingly, the new *T*_12_ test (inspired by *T*_*ME*_ but based on ***s***) has comparable power with *T*_*skato*_, but without the need to search for the ‘optimal’ *ρ*. Our simulation results here also confirm that the three existing hybrid tests, *T*_*skato*_, *T*_*Minp*_ and *T*_*Fisher*_, have similar performance, where *T*_*skato*_ and *T*_*Minp*_ perform more similarly with each other than with *T*_*Fisher*_.

**Figure 1:**
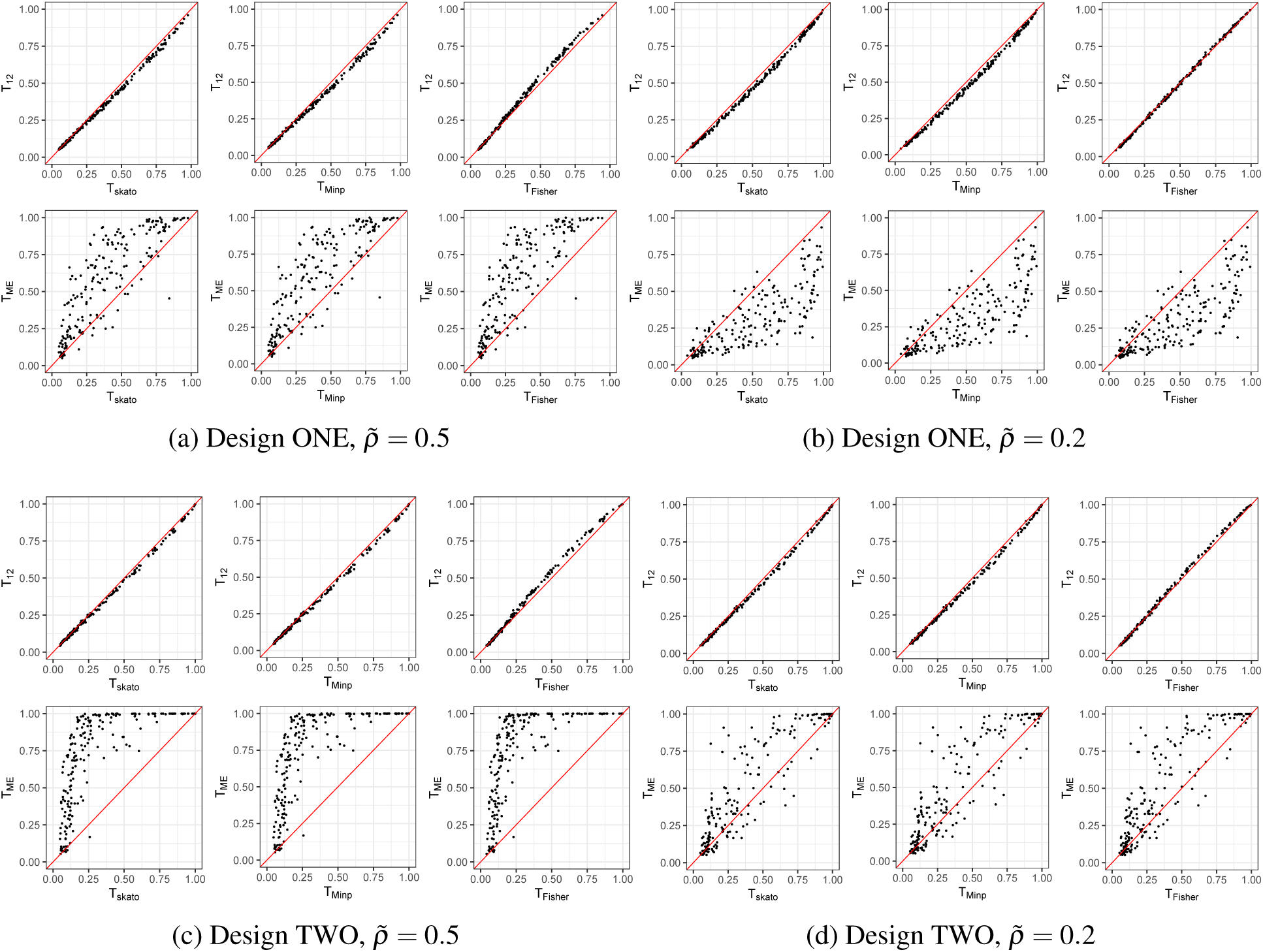
Power comparison under sparse alternatives based on design ONE and TWO, comparing *T*_12_ and *T*_*ME*_ with *T*_*Fisher*_, *T*_*Minp*_ **and** *T*_*skato*_. Empirical power of 200 alternative models are shown. For each model, *J* = 100 and the proportion of the causal variants varies from 1% to 10% randomly. See the main text for additional simulation details.

Figures 1(c) and 1(d) show that when ***s*** follows a mixed-effect model, the advantage of the proposed *T*_*ME*_ is enhanced. In this case, power of *T*_*ME*_ is considerably higher than the other tests even when 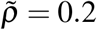 (Figures 1(d)).

### 3.3 Power with covariates

We now briefly study the effect of incorporating variant-specific additional information ***z*** = (*z*_1_, …, *z*_*J*_)^*T*^. As discussed before, although one may revise *w*_*j*_ to be proportional to *z* _*j*_ in additional to MAF, it is not immediately clear how to choose an ‘optimal’ weighting function. Thus, we only study the proposed *T*_*ME,cov*_, derived directly from the regression model (13), and we consider simulation design two only.

Without loss of generality, we assume *z* _*j*_ to be an indicator variable, for example indicating if the variant is non-synonymous (*z* _*j*_ = 1) or synonymous (*z* _*j*_ = 0). For causal variants we let *Pr*(*z*_*j*_ = 1) = 0.5, and for non-causal variants *Pr*(*z*_*j*_ = 1) = 0. We consider both the case of informative ***z*** (*θ* ≠ 0 in model (13) and *θ* ∼ Unif (1, 4)) and the case of uninformative ***z*** (*θ* = 0). Because of the additional information available from ***z***, we draw *β* from Unif (0.1, 1) and choose *τ*^2^ = 0. We also assumed *P*_*c*_ = 0.1 and *P*_*d*_ ∼ Unif (0.5, 1) for this set of simulations.

Figure 2 show that, as expected, there can be substantial power gain when incorporating informative covariate information (the left plot), at the cost of slightly reduced power when ***z*** is uninformative (the right plot).

**Figure 2:**
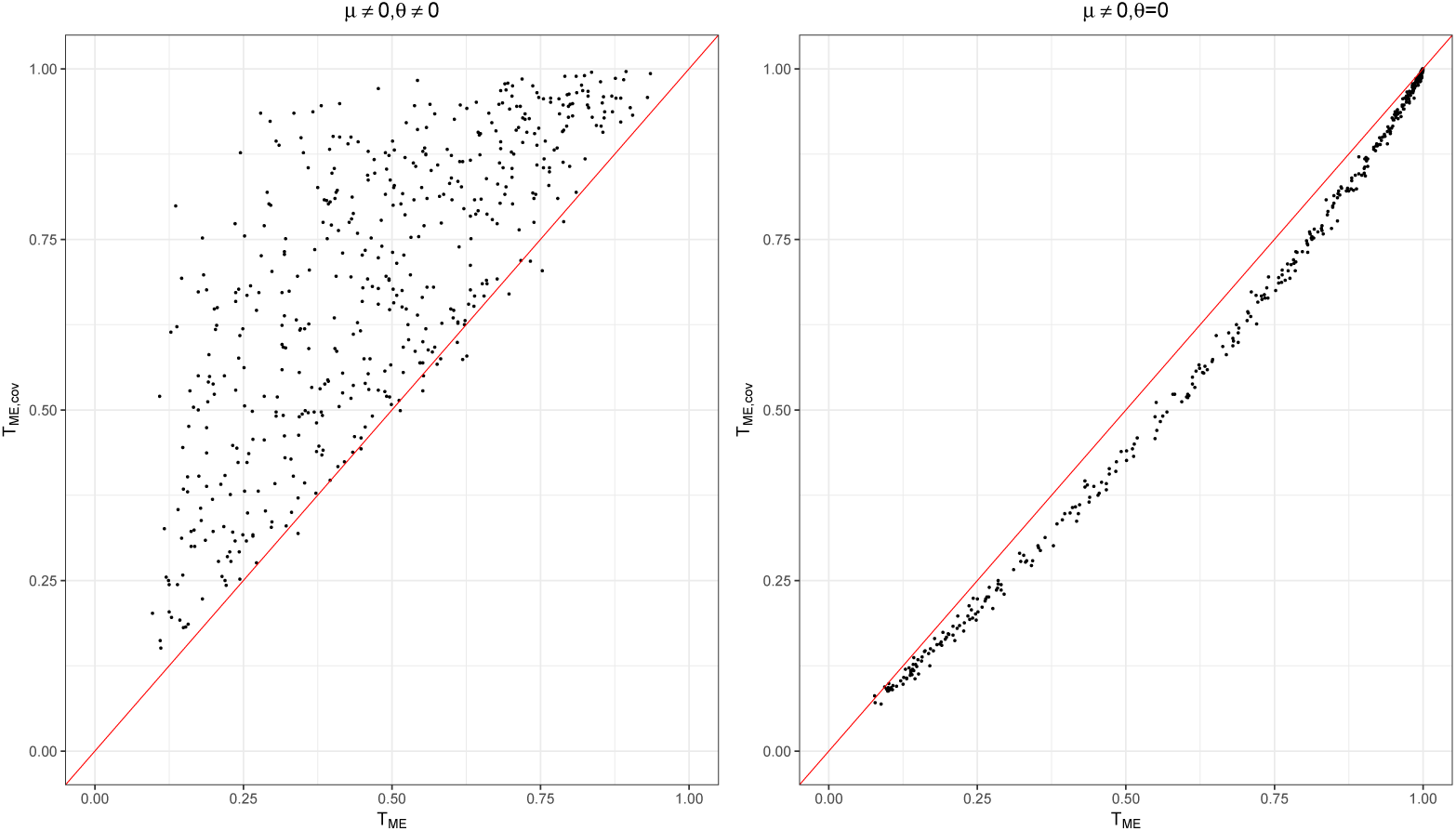
Power comparison under sparse alternatives based on design TWO, comparing *T*_*ME,cov*_ with *T*_*ME*_. The left plot is for the case of informative covariate (*θ* ≠ 0), and the right plot is for uninformative covariate (*θ* = 0). Empirical power of 500 alternative models are shown. For each model, *J* = 100 and the proportion of the causal variants is 10%, and 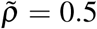. See the main text for additional simulation details.

## 4 Applications

In this section, we examine eight test statistics including *T*_*ME,cov*_, *T*_*ME*_ and *T*_12_, and the existing methods, *T*_*skato*_, *T*_*Minp*_, *T*_*Fisher*_, as well as *T*_1_ and *T*_2_ for completeness through two data applications. In the implementation of *T*_*ME,cov*_, we use variants being non-synonymous or synonymous, annotated using the UCSC genome browser at https://genome.ucsc.edu/, as the variant-specific information.

The first application highlights the advantage of the proposed *T*_*ME*_ in the presence of high or moderately high correlation between variants, and it also demonstrates that the method is not limited to analyses of rare variants. The second application revisits the genetic analysis workshop 17 (GAW17) rare variants data previously studied by Derkach, Lawless & Sun (2014). This application reveals the benefit of incorporating covariate information using *T*_*ME,cov*_. For both applications, we used method of Rothman (2012) to achieve a positive definite estimate for the covariance matrix **Σ**_0_.

### 4.1 Cystic Fibrosis (CF) data - common variants

Cystic Fibrosis is a life-limiting genetic condition for which lung function is a primary co-morbidity of interest. To *indirectly* study gene-environment interactions, Soave et al. (2015) proposed a joint location-scale (JLS) test and applied it to lung function measures in CF individuals, *n* = 1, 409 from a Canadian sample and *n* = 1, 232 from a French sample. They discovered and replicated the significance of the SLC9A3 complex set (35 common variants from four genes) based on the JLS test. However, the association evidence appears to come from the scale (interaction) component. For the traditional location (mean) test based on *T*_2_, the SLC9A3 complex set was only significant in the Canadian sample but not replicated in the French sample. Here we exam the performance of the eight tests applied to the French sample. For a fair comparison, we use ***w*** = 1, ***A*** = ***I*** and ***R*** = ***I*** for all corresponding tests, and we obtain empirical p-values for all tests based on 10^4^ permutation replicates.

Results in Table 3 show that only some of the genes appear to be truly associated with CF lung function. For SLC9A3, all tests have suggestive evidence with *T*_1_ having p-value < 0.05. For SLC9A3R1, benefiting from the correlation structure (Supplementary Material) the proposed *T*_*ME*_ and *T*_*ME,cov*_, which use 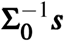 instead of ***s***, are significant. When jointly analyzing all four genes in the SLC9A3 complex set, none of the tests is statistically significant but *T*_*ME*_ has the smallest p-value. A larger sample is needed to make a definitive conclusion of true association. The covariate information (non-synonymous vs. synonymous) appear not to be informative here, but the performance of *T*_*ME,cov*_ is similar to that of *T*_*ME*_.

**Table 3:** Empirical p-values of the tests in the CF data application.

| Gene | $J$ | Empirical p-values | | | | | | | |
| --- | --- | --- | --- | --- | --- | --- | --- | --- | --- |
| | | $T_1$ | $T_2$ | $T_{skato}$ | $T_{Minp}$ | $T_{Fisher}$ | $T_{12}$ | $T_{ME}$ | $T_{ME,cov}$ |
| SLC9A3 | 7 | **0.0443** | 0.1198 | 0.079 | 0.0783 | 0.0541 | 0.0718 | 0.0886 | 0.0860 |
| EZR | 10 | 0.2192 | 0.6856 | 0.3241 | 0.3209 | 0.3897 | 0.239 | 0.3473 | 0.3474 |
| SLC9A3R2 | 10 | 0.804 | 0.5951 | 0.6965 | 0.6984 | 0.709 | 0.9037 | 0.9683 | 0.8791 |
| SLC9A3R1 | 8 | 0.0999 | 0.1103 | 0.1471 | 0.1656 | 0.0846 | 0.1042 | **0.0243** | **0.0243** |
| 4-gene jointly | 35 | 0.8372 | 0.3079 | 0.4749 | 0.4738 | 0.5671 | 0.9261 | 0.1710 | 0.1709 |

### 4.2 The Genetic Analysis Workshop 17 (GAW17) data - rare variants

Here we apply the method to the GAW17 data provided by the 1000 Genomes Project (Consortium et al. 2010; Almasy et al. 2011), focusing on the simulated quantitative trait Q2. The phenotype Q2 is influenced by 72 variants in 13 genes but not by environmental factors, and the genotypes of these variants are obtained from a ‘mini-exome’ next-generation sequencing experiment (Consortium et al. 2010). Available to us are 200 replicates, simulated based on a true phenotype-genotype association model determined by the GAW17 study group (Almasy et al. 2011) but blinded to this analysis. We consider *n* = 321 unrelated Asian samples and use only variants with MAF less than 0.05. The description of the variants is provided in Table 4. Among the 13 genes, GCKR is not analyzed since only one variant remained after variant screening. VNN1 does not have any causal rare variants but is kept for negative control.

**Table 4:** Empirical power of the tests in the GAW17 data application.

| Gene | $J_C$ | $J_N$ | Empirical Power | | | | | | | |
| --- | --- | --- | --- | --- | --- | --- | --- | --- | --- | --- |
| | | | $T_1$ | $T_2$ | $T_{skato}$ | $T_{Minp}$ | $T_{Fisher}$ | $T_{12}$ | $T_{ME}$ | $T_{ME,cov}$ |
| 7 genes for which the maximum power is 10% or more |  |  |  |  |  |  |  |  |  |  |
| SIRT1 | 4 | 7 | 0.44 | 0.385 | 0.455 | 0.43 | 0.495 | **0.5** | 0.285 | 0.285 |
| BCHE | 6 | 9 | 0.29 | 0.39 | 0.405 | 0.39 | 0.435 | 0.445 | **0.46** | 0.41 |
| PDGFD | 3 | 5 | 0.295 | 0.385 | 0.385 | 0.38 | 0.425 | **0.45** | 0.31 | 0.26 |
| SREBF1 | 4 | 5 | **0.495** | 0.25 | 0.440 | 0.440 | 0.445 | 0.400 | 0.405 | 0.355 |
| RARB | 1 | 5 | 0.06 | 0.135 | 0.095 | 0.110 | 0.085 | 0.090 | 0.100 | **0.215** |
| PLAT | 4 | 7 | 0.13 | 0.125 | 0.105 | 0.100 | 0.135 | 0.12 | 0.13 | **0.165** |
| VLDLR | 4 | 3 | 0.11 | 0.085 | 0.100 | 0.095 | 0.115 | 0.115 | 0.110 | **0.16** |
| 7-gene jointly | 26 | 41 | 0.935 | 0.765 | 0.94 | 0.935 | **0.945** | 0.94 | 0.765 | 0.765 |
| 4 genes for which the maximum power is 10% or less |  |  |  |  |  |  |  |  |  |  |
| VNN3 | 2 | 2 | 0.035 | 0.04 | 0.035 | 0.035 | 0.04 | 0.04 | 0.04 | 0.035 |
| INSIG1 | 3 | 1 | 0.05 | 0.03 | 0.03 | 0.03 | 0.035 | 0.035 | 0.025 | 0.015 |
| LPL | 1 | 3 | 0.03 | 0.065 | 0.035 | 0.035 | 0.04 | 0.035 | 0.035 | 0.050 |
| VWF | 1 | 3 | 0.025 | 0.01 | 0.01 | 0.01 | 0.015 | 0.01 | 0.04 | 0.045 |
| 4-gene jointly | 7 | 9 | 0.075 | 0.01 | 0.06 | 0.06 | 0.055 | 0.045 | 0.055 | 0.045 |
| 1 gene for which there is no rare causal variants, used as a negative control |  |  |  |  |  |  |  |  |  |  |
| VNN1 | 0 | 3 | 0.015 | 0.045 | 0.045 | 0.045 | 0.035 | 0.04 | 0.04 | 0.045 |

For each of the 200 alternative replicates, we calculate the empirical p-values (based on 10^4^ permutation replicates) for the eight test statistics. For each test, the power for *α* = 0.05 is estimated as the proportion of the 200 replicates for which the empirical p-values ≤ 0.05. We separate the 11 genes into three categories based on power as in Derkach, Lawless & Sun (2014) (which examined *T*_1_ and *T*_2_) and Derkach, Lawless & Sun (2013) (which examined *T*_1_, *T*_2_, *T*_*Minp*_, and *T*_*Fisher*_), and we also jointly analyze all genes within each category.

In this application, because the correlation is weak among variants (Supplementary Material), we anticipate that methods relies on ***s*** will have better power than those based on 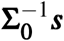. Indeed, results in Table 4 show that *T*_12_ has better performance than *T*_*ME*_. However, this application clearly demonstrates the potential of incorporating informative covariates. For example, the power of analyzing RARB, PLAT and VLDLR is higher using *T*_*ME,cov*_, at the cost of slightly reduced performance for other genes when the included covariate is presumably not (detectably) informative. Interestingly, *T*_*Fisher*_ has comparable performance as *T*_12_, and both outperform *T*_*skato*_ in almost all cases. Although the individual *T*_1_ and *T*_2_ tests may have the highest power for certain genes, the robustness of the hybrid tests is evident based on the overall performance exhibited in Table 4.

## 5 Discussion

In this paper, we considered a summary statistics-based regression framework to jointly analyze a set of *J* variants. As delineated in Table 1, the proposed approach is flexible and adaptive. The Empirical Power score test derived from the fixed-effect model, *T*_*FE*_, unifies the linear class of tests (also known as the burden tests), *T*_1_, while *T*_*RE*_ from the random-effect model connects the quadratic class or variance component tests, *T*_2_. Further, the score test derived from the random-effect model offers a new hybrid test, *T*_*ME*_, that naturally aggregates information from *T*_*FE*_ and *T*_*RE*_.

It is worth emphasizing three notable differences between the proposed *T*_*ME*_ and the well-known *T*_*skato*_ (Lee, Wu & Lin, 2012). First, the proposed framework aggregates evidence across *J* variants based on 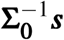, a precision matrix-based transformation of the score vector, that can increase power for detecting sparse alternatives (Fan, Jin & Yao, 2013; Cai, Liu & Xia, 2014). Secondly, unlike *T*_*skato*_, *T*_*ME*_ naturally aggregates the information from the linear and quadratic types of tests, based on a regression model *without* searching for the ‘optimal’ weight. Lastly, when additional variant-specific information is available, it is straightforward to derive *T*_*ME,cov*_ that accounts for covariate effects. We have demonstrated these features of the proposed method both analytically and empirically.

To exploit the assumption of signal sparsity, various supremum-type tests have been proposed including the generalized higher criticism (Wu et al., 2014; Barnett, Mukherjee & Lin, 2017) for sparse signals, and most recently the generalized Berk-Jones statistic (Sun & Lin, 2019) for moderately sparse signals. These methods, tailored for common variants, are not easy to adjust for additional variant-specific information.

The proposed set-based testing framework is a general one, and it can be used for other settings such as pleiotropy studies of multiple phenotypes, where the analtyical unit is each of the phenotypes. In that context, Liu & Lin (2018b) also proposed a summary statistics-based linear mixed-effect regression model, but they focused on the special case of ***w*** = 1 and ***R*** = ***I***. In addition, Liu & Lin (2018b) derived two score test statistics, respectively, for testing *µ* = 0 and *τ*^2^ = 0 *separately*, then considered different ways to combine the evidence including SKAT-O type of statistics. In contrast, we derive *T*_*ME*_ from testing *µ* = 0 and *τ*^2^ = 0 *jointly*, and the weighting factors are inherently justified based on the regression model. We also study the asymptotic properties of the proposed tests under the alternatives as well as empirical property under sparse alternative.

The proposed method can also be used for the study of polygenic risk score (PRS) (Purcell et al., 2009), and the connection between PRS and *T*_1_ has been noted by Pan, Chen & Wei (2015). In principle, *T*_*ME*_ can overcome the poor statistical efficiency of *T*_1_, but accurate estimation of large precision matrices can be challenging and requires special considerations (Fan, Liao & Liu, 2016). The link between *T*_1_ and PrediXcan (Gamazon et al., 2015) and TWAS (Gusev et al., 2016) for association and tissue-specific gene-expression data integration has also been noted (Xu et al. 2017). The performance of *T*_*ME*_ in this setting and comparison with other concurrently developed newer methods are of our future research interest.

Fix-, random- and mixed-effect models for summary statistics have been studied for meta-analyses of GWAS (Han & Eskin, 2011). In that context, a likelihood ratio test was implemented for a mixed-effect model, and the resulting test is also known as the new RE meta-analysis. The original test of Han & Eskin (2011) was designed for meta-analyses of independent studies, and a modified procedure has since been developed by Lee, Eskin & Han (2017) to account for correlations between studies but without adjusting for covariate effects. Comparisons between the two approaches for meta-analyses and other studies warrant future investigations.

## Acknowledgements

We thank Dr. Lisa J. Strug and her lab, and Dr. Harriet Corvol for providing the cystic fibrosis application data, and we thank the Genetic Analysis Workshop 17 (GAW17) committee and the 1000 Genomes Project for providing the GAW17 application data. We would also like to thank Dr. Andriy Derkach for helpful discussions. YZ is a trainee of the CIHR STAGE (Strategic Training in Advanced Genetic Epidemiology) training program at the University of Toronto. This research is funded by the Natural Sciences and Engineering Research Council of Canada (NSERC, RGPIN-04934 and RGPAS-522594), the Canadian Institutes of Health Research (CIHR, MOP-310732), and the University of Toronto McLaughlin Centre Accelerator Grants in Genomic Medicine (MC-2019-15).

